## Supporting Information for "Decoding reveals the neural representation of perceived and imagined musical sounds"

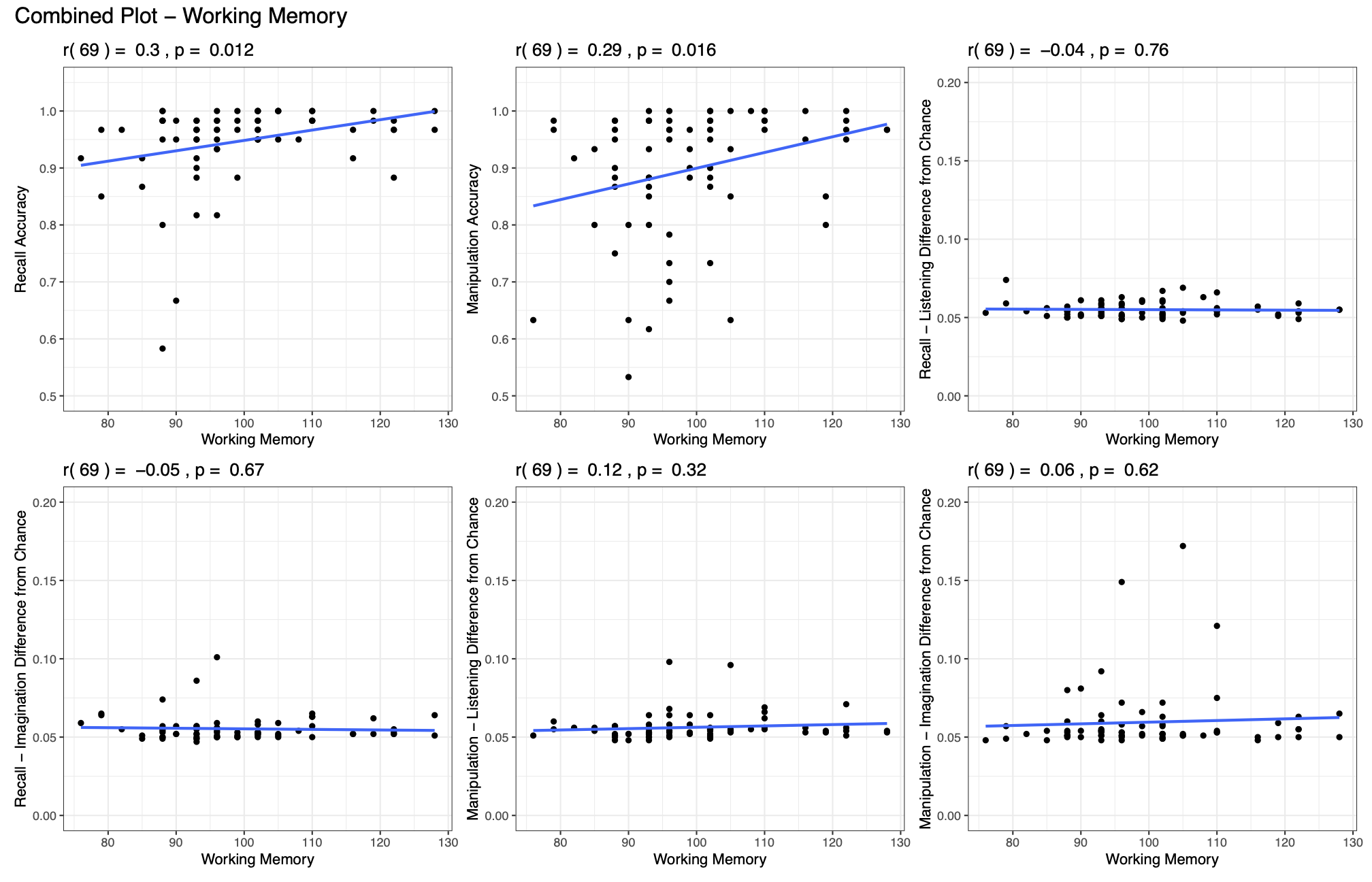

**Figure S1.** The relationship between working memory and recall and manipulation accuracy. We report Pearson correlations between working memory scores, calculated using the Wechsler Adult Intelligence Scale (WAIS), and recall and manipulation accuracy scores for each subject. Results show a positive correlation between working memory and both recall and manipulation accuracy. This relationship suggests that general working memory skills are beneficial for the retention and manipulation of (non-verbal) auditory objects in the context of an active imagery task.

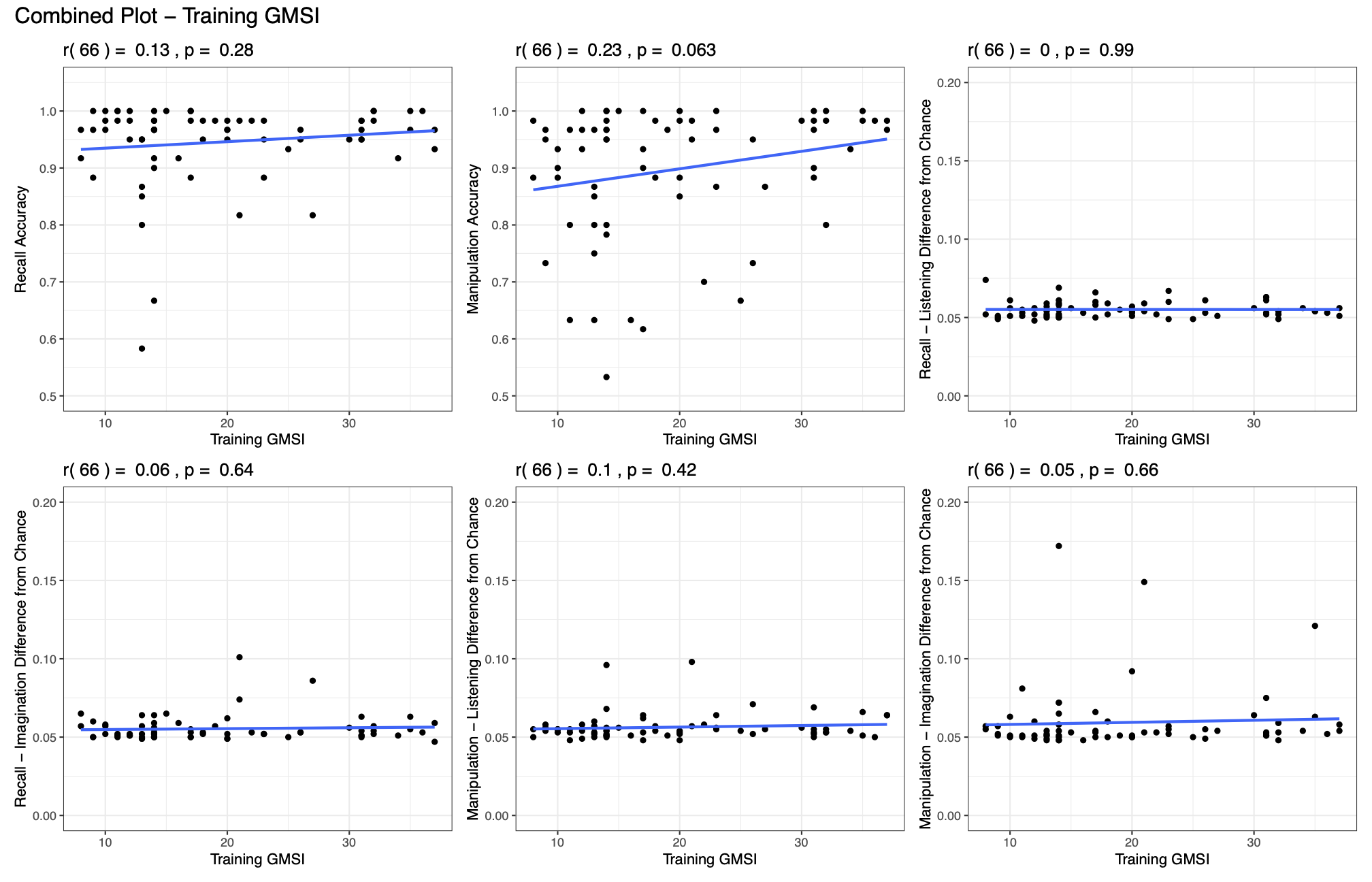

**Figure S2.** The relationship between music training (training subscale from the Goldsmiths Musical Sophistication Index - GMSI) and recall and manipulation accuracy. We report Pearson correlations between training GMSI scores, and recall and manipulation accuracy scores for each participant. Results show small positive correlations between music training and both recall and manipulation accuracy which, nonetheless, were statistically non-significant.

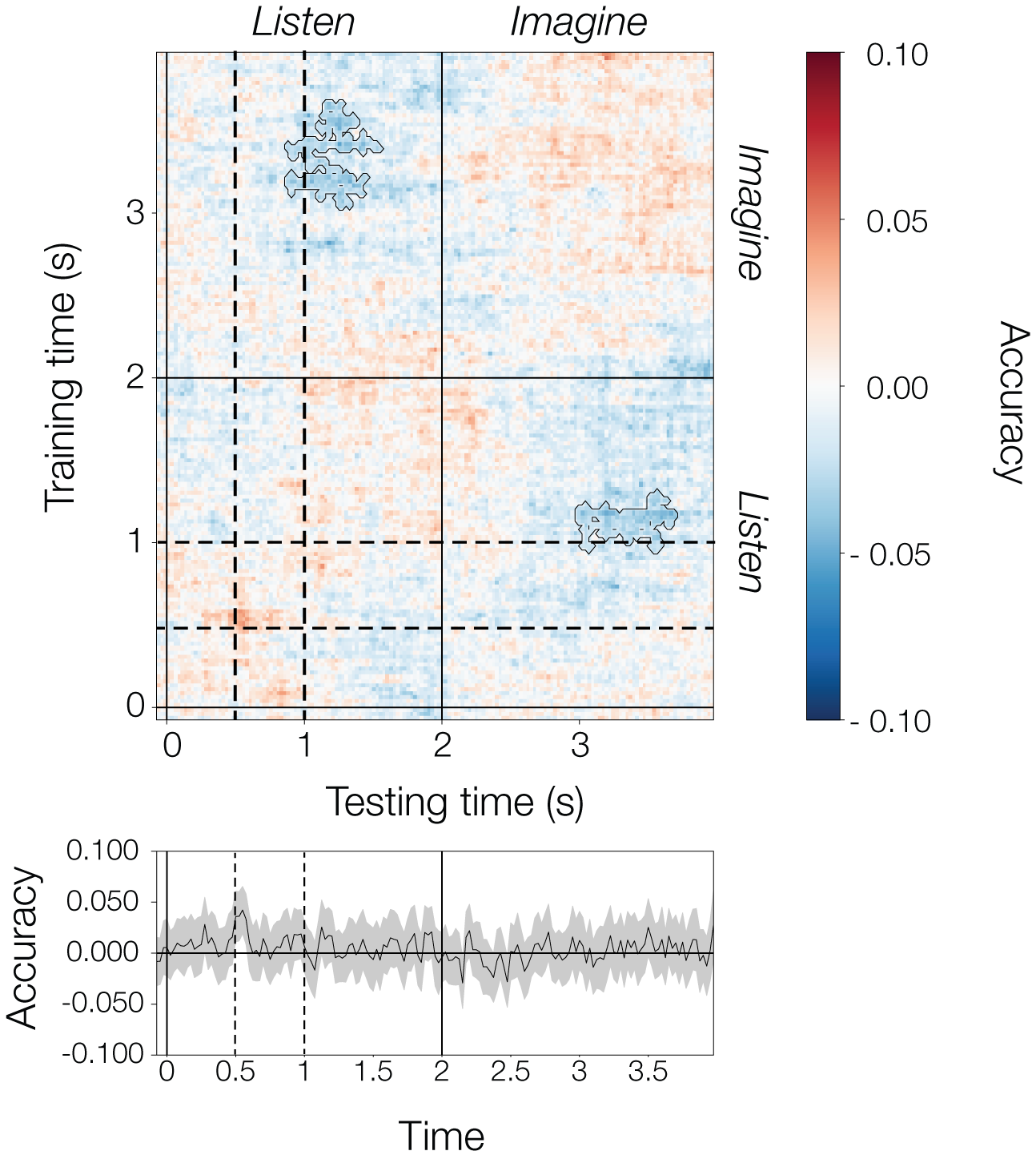

**Figure S3.** Difference between manipulation and recall for within-condition testing. Note how significant below-chance accuracy (contours) emerges when training on listening and testing on imagination and vice-versa. Bottom time-course depicts accuracy differences at the diagonal. Dashed lines mark the onset of the second (0.5s) and third (1s) sounds of the melodies.

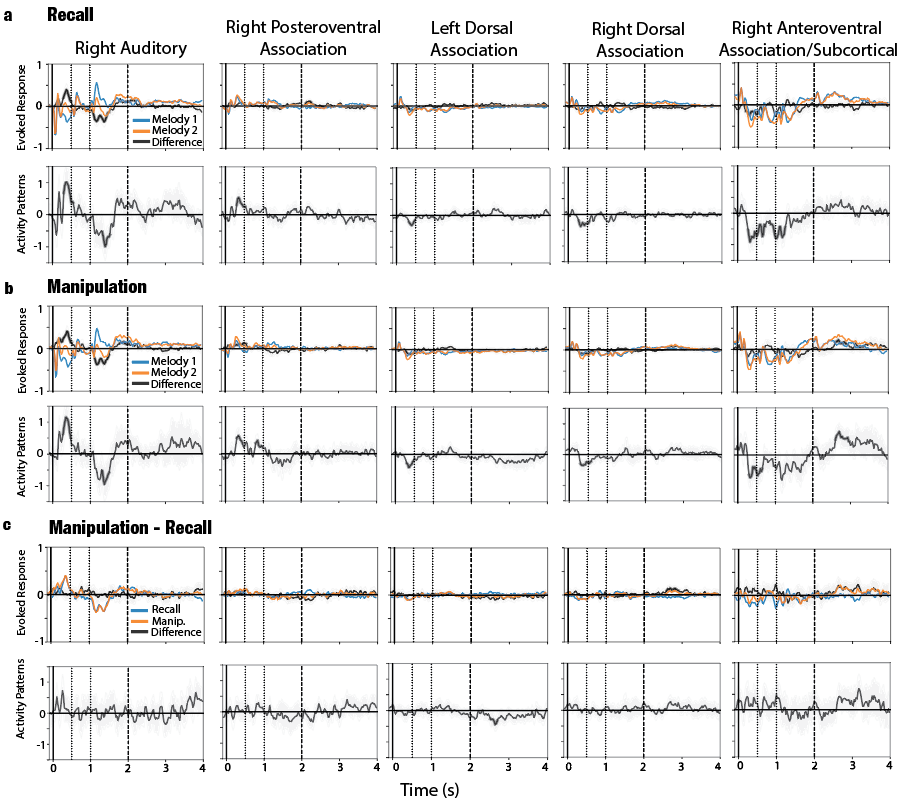

**Figure S4.** Time course of source-level evoked responses and decoding patterns for five different regions of interest, according to the Desikan-Killiany parcellation (1): ***Right Auditory*** (ctx-rh-superiortemporal, ctx-rh-bankssts, ctx-rh-transversetemporal), ***Right Posteroventral Association*** (ctx-rh-fusiform, ctx-rh-inferiortemporal, Right-Hippocampus, ctx-rh-parahippocampal', ctx-rh-isthmuscingulate, ctx-rh-precuneus,ctx-rh-lingual), ***Left Dorsal Association*** (ctx-lh-rostralmiddlefrontal, ctx-lh-superiorfrontal, ctx-lh-parstriangularis, ctx-lh-parsopercularis, ctx-lh-parsorbitalis, ctx-lh-insula), ***Right Dorsal Association*** (ctx-rh-rostralmiddlefrontal, ctx-rh-superiorfrontal, ctx-rh-parstriangularis, ctx-rh-parsopercularis, ctx-rh-parsorbitalis, ctx-rh-insula), and ***Right Anteroventral Association/Subcortical*** (ctx-rh-medialorbitofrontal, ctx-rh-lateralorbitofrontal, Right-Accumbens-area, Right-Caudate, Right-Putamen, Right-Pallidum, Right-Thalamus-Proper, ctx-rh-rostralanteriorcingulate). Bold segments indicate significant periods.

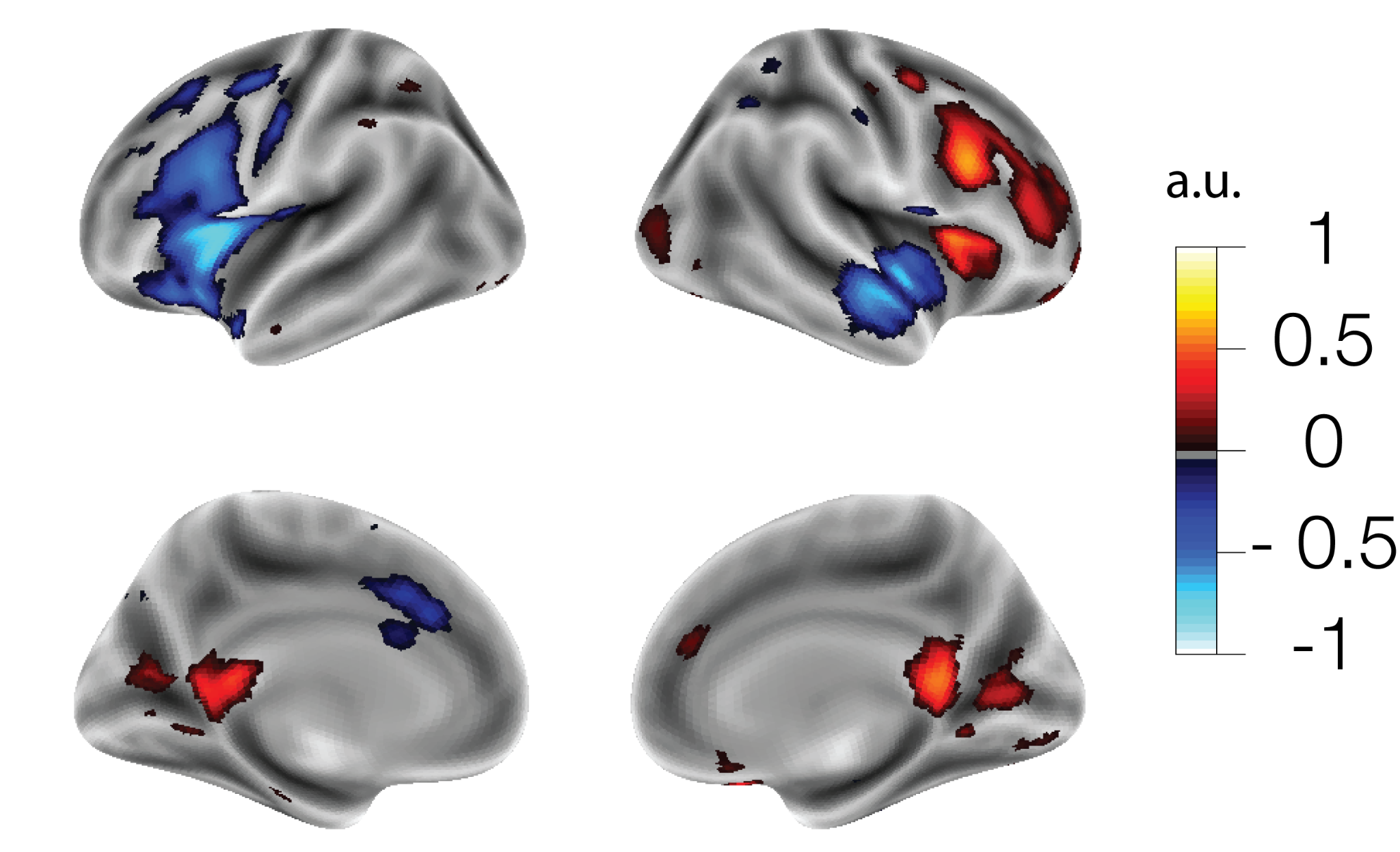

**Figure S5.** Uncorrected (t > 1.99) significant differences between manipulation and recall for the imagination period (2s - 4s). Note the bilateral frontal and medial parietal differences.

**Exploratory analysis: Relationship of neural decoding accuracy with behavioral measures**

For completeness, in exploratory analyses, we tested whether neural decoding accuracy was related to behavioral accuracy, musical training or vividness ratings. To this purpose, we took the mean of neural decoding accuracy scores in the generalization matrices (**Fig. 2**) for the listening (0s - 2s) and imagination (2s - 4s) periods separately (i.e., the left-bottom and top-right quadrants). Since below-chance and above-chance accuracy values were present, we derived an unsigned neural discrimination measure (**ND**) by subtracting 0.5 (i.e., the chance level) and computing the absolute value before taking the mean of the corresponding period. ND represents the information in the neural signal that drives above or below-chance decoding. In the manipulation condition, ND was significantly correlated with behavioral accuracy for the listening period (r = .24, p = .045) and marginally for the imagination period (r = .22, p = .052), before multiple comparisons corrections (**Table S1**). No other significant associations were found.

**Table S1.** Pearson’s correlation coefficients (***r***), and corresponding *p*-values with (***p_FDR_***) and without (***p***) false-discovery-rate correction, between neural discrimination (**ND**) and three variables of interest: Behavioral accuracy, vividness ratings and music training (GMSI scores).

| **Condition / Period** | **Behavioral Accuracy** | | | | **Vividness Ratings** | | | | **Music Training (GMSI)** | | |
| --- | --- | --- | --- | --- | --- | --- | --- | --- | --- | --- | --- |
|  | ***r*** | ***p*** | ***p_FDR_*** | ***r*** | | ***p*** | ***p_FDR_*** | ***r*** | | ***p*** | ***p_FDR_*** |
| Recall / listening | -.07 | .553 | .885 | .08 | | .523 | .885 | 0 | | .989 | .989 |
| Recall / imagination | -.15 | .224 | .674 | .16 | | .198 | .673 | .06 | | .639 | .885 |
| Manipulation / listening | **.24** | **.045** | .315 | -.01 | | .924 | .989 | .1 | | .425 | .885 |
| Manipulation / imagination | .23 | .052 | .315 | -.04 | | .763 | .916 | .05 | | .664 | .885 |

In addition, we explored whether the likelihood of a correct answer in the task (and corresponding response times), was related to the likelihood of correct neural decoding in the listening and imagination periods, at the single-trial level. We used logistic mixed-effects models (R, lme4) (2, 3) to evaluate whether participants’ target melody identification predicted correct melody discrimination from neural activity at each time point of the trial. We did this analysis separately for the recall and manipulation blocks, and for the listening and imagination periods. We included participant as random effect for both intercept and slopes. We obtained *p*-values for the relevant coefficients using the Satterthwaite approximation of degrees of freedom (4). The same analysis was done to predict neural accuracy from response times, as measured from the beginning of the test melody. No significant relationships were detected in these analyses, likely due to the ceiling effect in behavioral performance (**Fig. S6** and **S7)**.
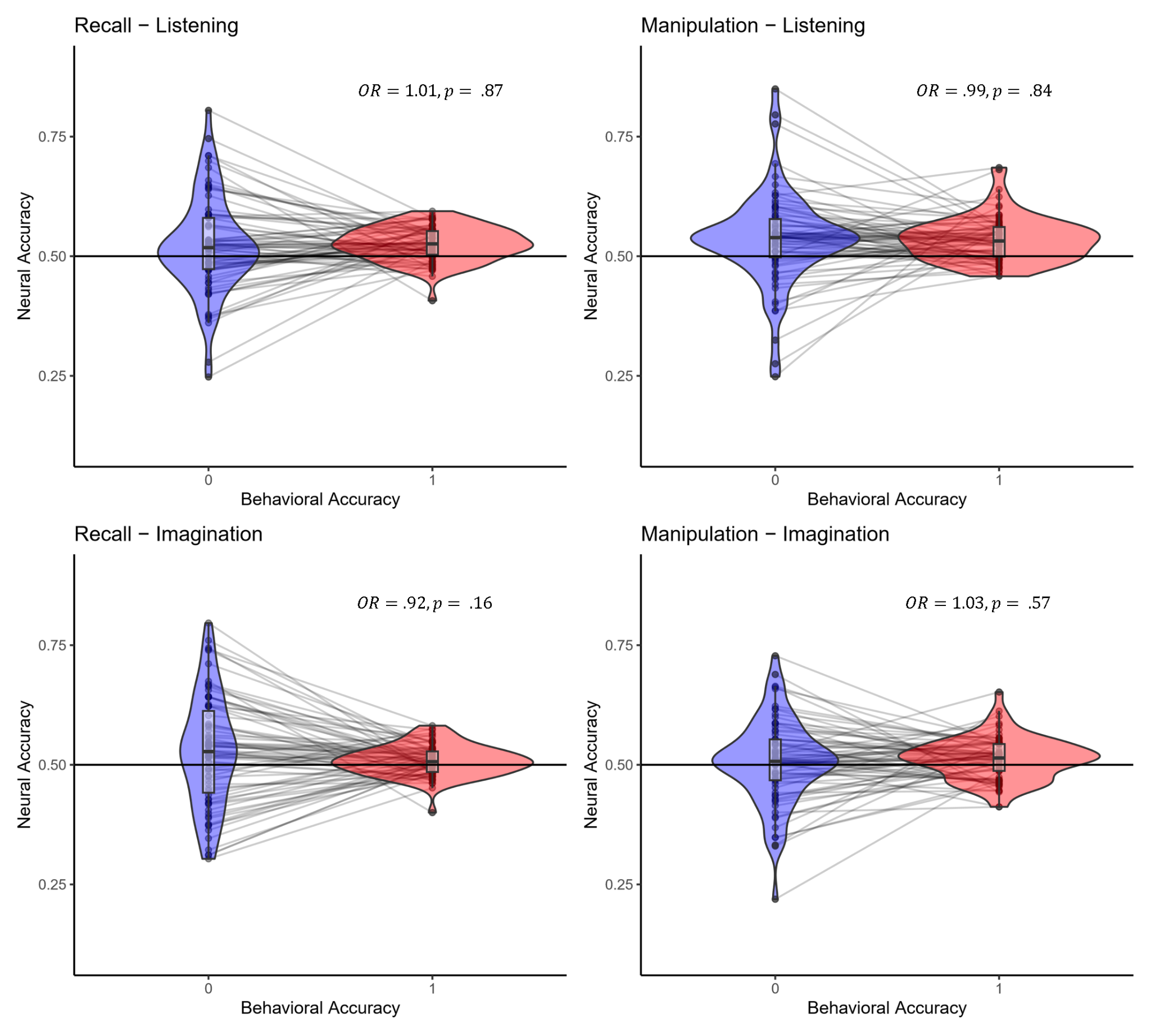

**Figure S6.** Relationship between neural decoding accuracy and behavioral performance at the single trial level. Using logistic regression (i.e., generalized mixed-effects models) we evaluated how well behavioral accuracy (correct=1, incorrect=0) predicted neural decoding of melody identity across all trials, participants and timepoints for the two conditions (recall, manipulation) and two different periods (listening, imagination) separately. The models included participant as a random effect (both for intercept and slopes). Each dot represents a prediction from the model for a single subject. Boxplots depict the median and interquartile range. The larger variance for incorrect trials reflects their smaller number compared to correct trials. Note that some participants did not exhibit any incorrect trials, but the model still extrapolated neural accuracy coefficients for them.

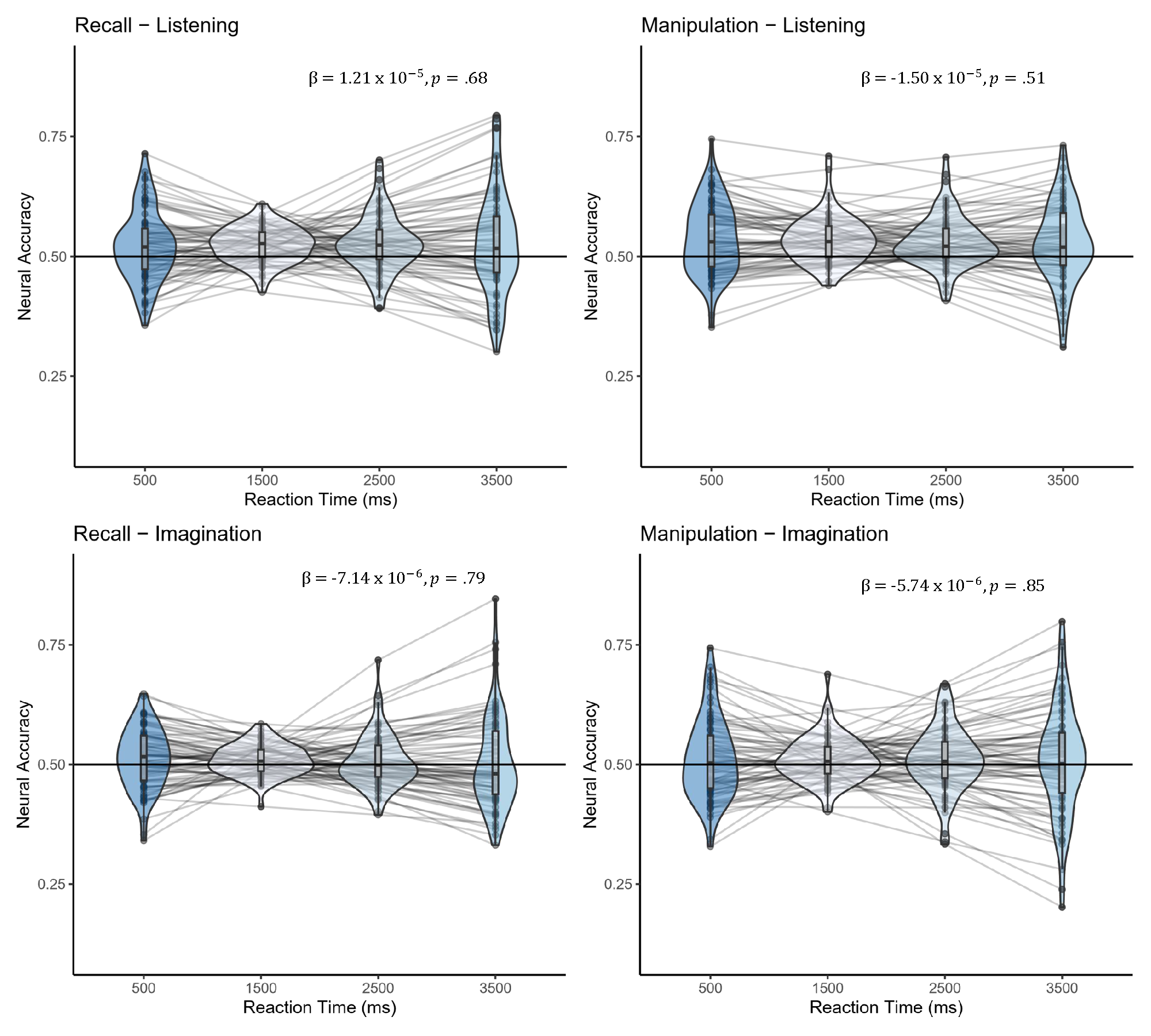

**Figure S7.** Relationship between neural decoding accuracy and reaction time at the single trial level. Using logistic regression (i.e., generalized mixed-effects models), we evaluated how well reaction time predicted neural decoding of melody identity across all trials, participants and time points for the two conditions (recall, manipulation) and two different periods (listening, imagination) separately. The models included participant as a random effect (both for intercept and slope). Each dot represents a prediction from the model for a single subject at different RTs (500ms, 1500ms, 2500ms, 3500ms). Boxplots depict the median and interquartile range.

**Figure S8.** Source localization of decoding patterns using inverse operators estimated with the data covariance of the listening (sounds 1, 2, and 3) and imagination period separately. Significant clusters are displayed.

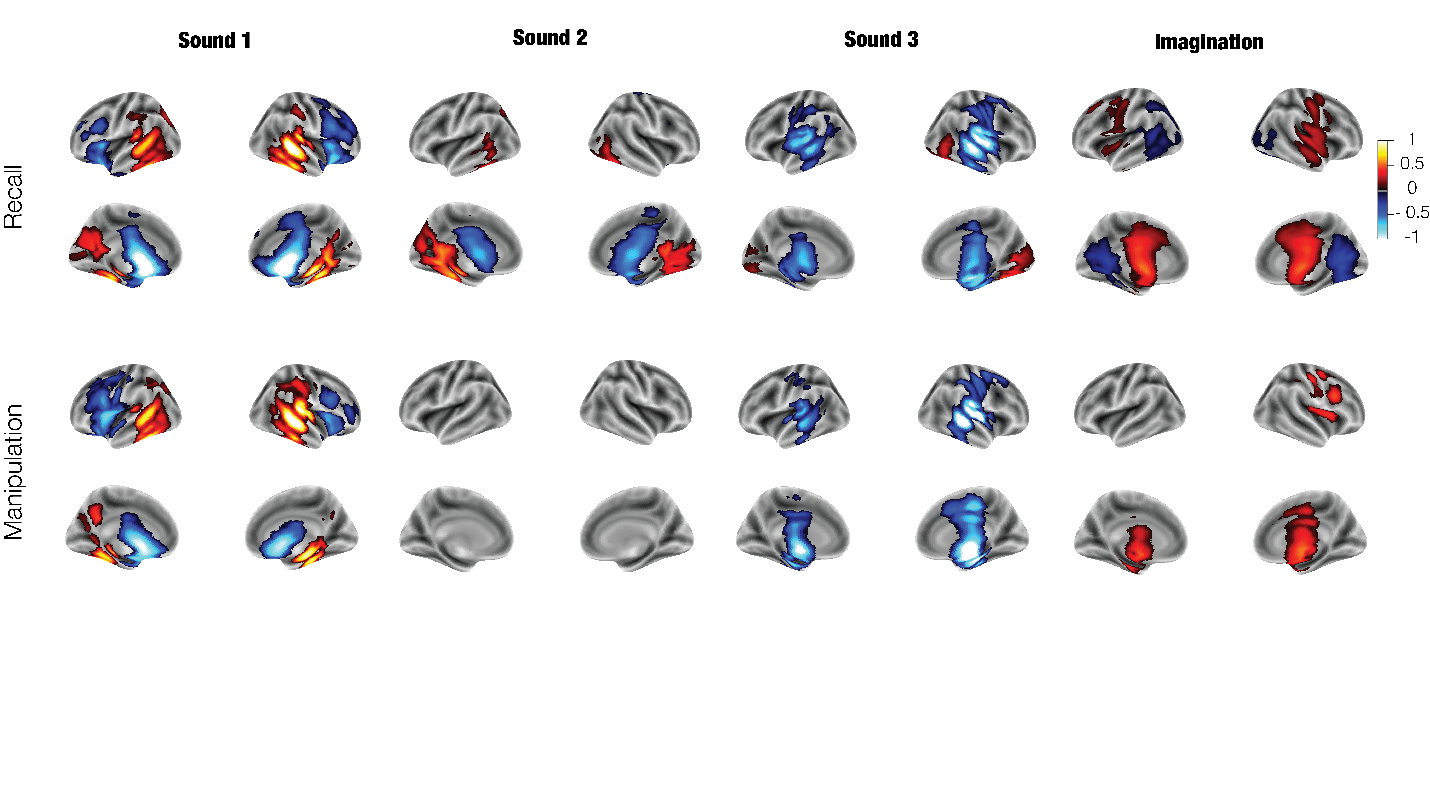

**Table S2.** Summary of all significant clusters found for the tests performed on time-generalized accuracy matrices.

| **condition** | **training peak time (s)** | **testing peak time (s)** | **cluster p-value** | **peak t-value** | **peak accuracy** |
| --- | --- | --- | --- | --- | --- |
| train on recall and test on recall | 0.2 | 0.2 | 0 | 7.13 | 0.573 |
|  | 1.23 | 1.23 | 0 | 7.949 | 0.589 |
|  | 2.33 | 2.28 | 0 | 6.863 | 0.547 |
|  | 3.15 | 3.1 | 0 | 5.555 | 0.543 |
|  | 1.38 | 0.53 | 0.007 | -2.185 | 0.485 |
|  | 0.18 | 1.42 | 0.008 | -2.109 | 0.486 |
| train on manipulation and test on manipulation | 1.2 | 1.2 | 0 | 9.675 | 0.602 |
|  | 3.35 | 3.65 | 0 | 4.542 | 0.549 |
|  | 1.18 | 0.5 | 0.016 | -2.019 | 0.486 |
|  | 0.53 | 1.9 | 0.014 | -2.017 | 0.485 |
| train on recall and test on manipulation | 0.2 | 0.18 | 0 | 7.767 | 0.569 |
|  | 1.2 | 1.2 | 0 | 7.927 | 0.577 |
|  | 1.4 | 0.18 | 0 | -2.174 | 0.488 |
|  | 0.08 | 1.42 | 0 | -2.069 | 0.486 |
|  | 3.18 | 2.78 | 0.048 | -2.061 | 0.485 |
|  | 3.3 | 2.98 | 0.011 | -2.018 | 0.485 |
|  | 3.35 | 3.78 | 0.034 | -2.01 | 0.484 |
| train on manipulation and test on recall | 0.18 | 0.18 | 0 | 9.901 | 0.58 |
|  | 1.2 | 1.2 | 0 | 8.008 | 0.574 |
|  | 1.5 | 0.35 | 0 | -2.06 | 0.487 |
|  | 0.38 | 1.63 | 0 | -2.047 | 0.485 |
|  | 2.9 | 2.92 | 0.011 | -2.056 | 0.487 |
|  | 3.28 | 3.52 | 0.018 | -2.039 | 0.485 |
| recall minus manipulation (within-condition testing) | 3.2 | 1.33 | 0.033 | -4.036 | -0.049 |
|  | 3.42 | 1.25 | 0.016 | -5.289 | -0.053 |
|  | 1.18 | 3.2 | 0.009 | -4.625 | -0.05 |
| test on recall (between minus within) | 2.3 | 2.28 | 0.03 | -5.762 | -0.058 |
|  | 3.38 | 3.45 | 0 | -5.563 | -0.067 |
| test on manipulation (between minus within) | 1.88 | 1.9 | 0.035 | -4.545 | -0.064 |
|  | 3.22 | 3.32 | 0 | -4.434 | -0.079 |

**Table S3.** Summary of all significant clusters found for the tests performed on patterns of activation at the source level. Coordinates of peak activity (x, y, z) are given in standard MNI space.

| **period or comparison** | **condition** | **cluster No** | **x** | **y** | **z** | **cluster p-value** | **peak t-value** | **peak activation** |
| --- | --- | --- | --- | --- | --- | --- | --- | --- |
| first sound | recall | 1 | 50 | -40 | -15 | 0.0002 | 6.19 | 0.91 |
|  |  | 2 | 0 | 5 | -5 | 0.0002 | -7.67 | -1.45 |
|  | manipulation | 1 | -65 | -55 | 0 | 0.0066 | 5.16 | 0.23 |
|  |  | 2 | 50 | -40 | -15 | 0.0048 | 5.31 | 0.83 |
|  |  | 3 | -20 | 5 | 15 | 0.0002 | -5.72 | -1.04 |
| second sound | recall | 1 | -30 | -40 | -25 | 0.0012 | 4.73 | 0.72 |
|  |  | 2 | 0 | -5 | 10 | 0.0046 | -4.98 | -0.88 |
|  | manipulation | 1 | -35 | -10 | 50 | 0.0014 | -4.61 | -0.42 |
| third sound | recall | 1 | 25 | -75 | -15 | 0.0214 | 4.23 | 0.39 |
|  |  | 2 | 0 | -15 | 5 | 0.0002 | -5.00 | -0.82 |
|  | manipulation | 1 | 25 | -15 | -5 | 0.0002 | -5.28 | -1.61 |
| imagination | recall | 1 | 0 | -10 | 25 | 0.0002 | 5.28 | 0.42 |
|  |  | 2 | 15 | -60 | 15 | 0.0008 | -4.67 | -0.31 |
|  | manipulation | 1 | 5 | -10 | 5 | 0.0008 | 4.64 | 0.48 |
|  | difference | 1 | -25 | 15 | 30 | 0.0328 | -3.53 | -0.48 |
| third minus first sound | recall | 1 | 0 | 25 | -20 | 0.0004 | 5.67 | 1.12 |
|  |  | 2 | 50 | -35 | -20 | 0.003 | -6.35 | -1.56 |
|  |  | 3 | -55 | -40 | -5 | 0.0172 | -4.78 | -1.23 |
|  | manipulation | 1 | -25 | 20 | 15 | 0.0038 | 4.16 | 0.87 |
|  |  | 2 | 55 | -35 | -5 | 0.0002 | -5.96 | -2.14 |
| imagination minus listening | recall | 1 | -5 | -10 | 5 | 0.0002 | 8.23 | 1.14 |
|  |  | 2 | -5 | -65 | 25 | 0.0002 | -6.5 | -0.44 |
|  | manipulation | 1 | 0 | -10 | 5 | 0.0002 | 6.64 | 0.93 |

**Table S4.** Peak t-value for each region in the **right** hemisphere that overlapped with significant clusters in each of the tested periods and conditions. The Desikan-Killiany parcellation was used to obtain anatomical labels.

| **Period/comparison** | **Condition** | **Cluster No** | **Label** | **t-value** |
| --- | --- | --- | --- | --- |
| First sound | recall | 1 | ctx-rh-inferiortemporal | 6.14 |
|  |  |  | ctx-rh-fusiform | 6.11 |
|  |  |  | ctx-rh-middletemporal | 5.32 |
|  |  |  | ctx-rh-bankssts | 5.29 |
|  |  |  | ctx-rh-superiortemporal | 4.66 |
|  |  |  | ctx-rh-parahippocampal | 4.46 |
|  |  |  | Right-Hippocampus | 4.16 |
|  |  |  | ctx-rh-inferiorparietal | 3.42 |
|  |  |  | ctx-rh-isthmuscingulate | 3.35 |
|  |  |  | ctx-rh-lingual | 3.32 |
|  |  |  | ctx-rh-transversetemporal | 3.16 |
|  |  |  | ctx-rh-supramarginal | 2.99 |
|  |  |  | ctx-rh-insula | 2.78 |
|  |  |  | ctx-rh-precuneus | 2.73 |
|  |  |  | Right-Thalamus-Proper | 2.61 |
|  |  |  | ctx-rh-postcentral | 2.58 |
|  |  | 2 | Right-Accumbens-area | -6.79 |
|  |  |  | ctx-rh-medialorbitofrontal | -6.47 |
|  |  |  | Right-Caudate | -6.26 |
|  |  |  | Right-Thalamus-Proper | -6.21 |
|  |  |  | ctx-rh-rostralanteriorcingulate | -5.8 |
|  |  |  | Right-Putamen | -5 |
|  |  |  | ctx-rh-lateralorbitofrontal | -4.63 |
|  |  |  | Right-Pallidum | -4.62 |
|  |  |  | ctx-rh-rostralmiddlefrontal | -4.3 |
|  |  |  | ctx-rh-insula | -4.26 |
|  |  |  | ctx-rh-parstriangularis | -4.01 |
|  |  |  | ctx-rh-parsopercularis | -3.88 |
|  |  |  | ctx-rh-parsorbitalis | -3.79 |
|  |  |  | Right-Amygdala | -3.56 |
|  |  |  | ctx-rh-caudalmiddlefrontal | -3.47 |
|  |  |  | ctx-rh-posteriorcingulate | -3.37 |
|  |  |  | ctx-rh-paracentral | -3.29 |
|  |  |  | ctx-rh-superiorfrontal | -3.29 |
|  |  |  | ctx-rh-superiortemporal | -3.14 |
|  |  |  | ctx-rh-entorhinal | -2.96 |
|  |  |  | ctx-rh-temporalpole | -2.77 |
|  |  |  | ctx-rh-precentral | -2.6 |
|  |  |  | Right-Hippocampus | -2.57 |
|  |  |  | ctx-rh-caudalanteriorcingulate | -2.54 |
|  | manipulation | 2 | ctx-rh-middletemporal | 5.23 |
|  |  |  | ctx-rh-inferiortemporal | 5.09 |
|  |  |  | ctx-rh-bankssts | 4.95 |
|  |  |  | ctx-rh-superiortemporal | 4.71 |
|  |  |  | ctx-rh-supramarginal | 4.49 |
|  |  |  | ctx-rh-inferiorparietal | 4.07 |
|  |  |  | ctx-rh-postcentral | 4.02 |
|  |  |  | ctx-rh-fusiform | 3.91 |
|  |  |  | ctx-rh-transversetemporal | 3.87 |
|  |  |  | ctx-rh-parahippocampal | 3.83 |
|  |  |  | Right-Hippocampus | 3.8 |
|  |  |  | ctx-rh-insula | 3.58 |
|  |  |  | ctx-rh-lingual | 2.71 |
|  |  |  | Right-Thalamus-Proper | 2.7 |
|  |  |  | ctx-rh-isthmuscingulate | 2.5 |
|  |  |  | ctx-rh-entorhinal | 2.39 |
|  |  |  | Right-Putamen | 2.27 |
|  |  | 3 | Right-Thalamus-Proper | -4.72 |
|  |  |  | Right-Accumbens-area | -4.65 |
|  |  |  | Right-Caudate | -4.38 |
|  |  |  | ctx-rh-medialorbitofrontal | -4.27 |
|  |  |  | ctx-rh-rostralanteriorcingulate | -3.78 |
|  |  |  | ctx-rh-superiorfrontal | -3.76 |
|  |  |  | ctx-rh-precentral | -3.48 |
|  |  |  | ctx-rh-insula | -3.42 |
|  |  |  | ctx-rh-lateralorbitofrontal | -3.37 |
|  |  |  | Right-Putamen | -3.34 |
|  |  |  | ctx-rh-rostralmiddlefrontal | -3.3 |
|  |  |  | ctx-rh-parstriangularis | -3.23 |
|  |  |  | ctx-rh-parsopercularis | -3.19 |
|  |  |  | ctx-rh-parsorbitalis | -2.96 |
|  |  |  | ctx-rh-caudalanteriorcingulate | -2.95 |
|  |  |  | Right-Pallidum | -2.95 |
|  |  |  | ctx-rh-caudalmiddlefrontal | -2.77 |
|  |  |  | ctx-rh-superiortemporal | -2.31 |
|  |  |  | ctx-lh-entorhinal | -2.3 |
| Second sound | recall | 1 | ctx-rh-isthmuscingulate | 3.72 |
|  |  |  | ctx-rh-lingual | 3.6 |
|  |  |  | ctx-rh-precuneus | 3.54 |
|  |  |  | ctx-rh-fusiform | 3.22 |
|  |  |  | ctx-rh-lateraloccipital | 3.17 |
|  |  |  | ctx-rh-pericalcarine | 3.14 |
|  |  |  | ctx-rh-cuneus | 2.8 |
|  |  |  | ctx-rh-inferiortemporal | 2.65 |
|  |  |  | ctx-rh-inferiorparietal | 2.57 |
|  |  |  | ctx-rh-parahippocampal | 2.54 |
|  |  |  | Right-Hippocampus | 2.35 |
|  |  | 2 | Right-Thalamus-Proper | -4.91 |
|  |  |  | Right-Caudate | -4.25 |
|  |  |  | Right-Amygdala | -3.73 |
|  |  |  | Right-Accumbens-area | -3.63 |
|  |  |  | ctx-rh-posteriorcingulate | -3.59 |
|  |  |  | Right-Putamen | -3.51 |
|  |  |  | Right-Pallidum | -3.51 |
|  |  |  | Right-Hippocampus | -3.33 |
|  |  |  | ctx-rh-entorhinal | -3.14 |
|  |  |  | ctx-rh-medialorbitofrontal | -3.01 |
|  |  |  | ctx-rh-lateralorbitofrontal | -2.8 |
|  |  |  | ctx-rh-paracentral | -2.71 |
|  |  |  | ctx-rh-rostralanteriorcingulate | -2.55 |
|  |  |  | ctx-rh-precentral | -2.53 |
|  |  |  | ctx-rh-precuneus | -2.51 |
|  |  |  | ctx-rh-caudalanteriorcingulate | -2.41 |
|  |  |  | ctx-rh-caudalmiddlefrontal | -2.35 |
|  |  |  | ctx-rh-insula | -2.15 |
|  |  |  | ctx-rh-isthmuscingulate | -2.03 |
|  | manipulation | 1 | ctx-rh-precentral | -3.96 |
|  |  |  | Right-Thalamus-Proper | -3.55 |
|  |  |  | ctx-rh-caudalmiddlefrontal | -3.35 |
|  |  |  | ctx-rh-parsopercularis | -3.27 |
|  |  |  | ctx-rh-insula | -3.1 |
|  |  |  | ctx-rh-rostralmiddlefrontal | -3.04 |
|  |  |  | Right-Putamen | -3.04 |
|  |  |  | ctx-rh-posteriorcingulate | -3.03 |
|  |  |  | ctx-rh-postcentral | -2.78 |
|  |  |  | ctx-rh-superiorfrontal | -2.74 |
|  |  |  | ctx-rh-caudalanteriorcingulate | -2.68 |
|  |  |  | ctx-rh-paracentral | -2.57 |
|  |  |  | Right-Pallidum | -2.4 |
|  |  |  | Right-Caudate | -2.37 |
|  |  |  | ctx-rh-supramarginal | -2.25 |
| Third sound | recall | 1 | ctx-rh-fusiform | 3.81 |
|  |  |  | ctx-rh-lingual | 3.73 |
|  |  |  | ctx-rh-pericalcarine | 3.69 |
|  |  |  | ctx-rh-lateraloccipital | 3.63 |
|  |  |  | ctx-rh-cuneus | 3.62 |
|  |  |  | ctx-rh-middletemporal | 2.67 |
|  |  |  | ctx-rh-inferiortemporal | 2.46 |
|  |  |  | ctx-rh-inferiorparietal | 2.33 |
|  |  | 2 | Right-Thalamus-Proper | -4.94 |
|  |  |  | ctx-rh-superiortemporal | -4.78 |
|  |  |  | ctx-rh-inferiortemporal | -4.69 |
|  |  |  | ctx-rh-postcentral | -4.59 |
|  |  |  | ctx-rh-middletemporal | -4.53 |
|  |  |  | Right-Pallidum | -4.49 |
|  |  |  | ctx-rh-supramarginal | -4.46 |
|  |  |  | ctx-rh-bankssts | -4.34 |
|  |  |  | ctx-rh-transversetemporal | -4.31 |
|  |  |  | ctx-rh-fusiform | -4.28 |
|  |  |  | Right-Putamen | -4.23 |
|  |  |  | Right-Amygdala | -4.13 |
|  |  |  | Right-Hippocampus | -4.07 |
|  |  |  | ctx-rh-precentral | -3.84 |
|  |  |  | ctx-rh-insula | -3.82 |
|  |  |  | ctx-rh-entorhinal | -3.76 |
|  |  |  | ctx-rh-parahippocampal | -3.59 |
|  |  |  | Right-Caudate | -3.24 |
|  |  |  | ctx-rh-posteriorcingulate | -2.92 |
|  |  |  | ctx-rh-paracentral | -2.61 |
|  |  |  | ctx-rh-superiorfrontal | -2.57 |
|  |  |  | ctx-rh-caudalmiddlefrontal | -2.52 |
|  |  |  | ctx-rh-parsopercularis | -2.45 |
|  |  |  | Right-Accumbens-area | -2.29 |
|  |  |  | ctx-rh-rostralmiddlefrontal | -2.01 |
|  | manipulation | 1 | Right-Pallidum | -5.28 |
|  |  |  | Right-Putamen | -5.22 |
|  |  |  | Right-Thalamus-Proper | -4.76 |
|  |  |  | ctx-rh-precentral | -4.71 |
|  |  |  | Right-Caudate | -4.6 |
|  |  |  | ctx-rh-supramarginal | -4.58 |
|  |  |  | Right-Amygdala | -4.51 |
|  |  |  | Right-Hippocampus | -4.44 |
|  |  |  | ctx-rh-postcentral | -4.43 |
|  |  |  | ctx-rh-middletemporal | -4.36 |
|  |  |  | ctx-rh-posteriorcingulate | -4.32 |
|  |  |  | ctx-rh-superiortemporal | -4.28 |
|  |  |  | ctx-rh-paracentral | -4.08 |
|  |  |  | ctx-rh-bankssts | -4.03 |
|  |  |  | ctx-rh-insula | -3.9 |
|  |  |  | ctx-rh-superiorfrontal | -3.84 |
|  |  |  | ctx-rh-transversetemporal | -3.6 |
|  |  |  | ctx-rh-caudalanteriorcingulate | -3.38 |
|  |  |  | ctx-rh-caudalmiddlefrontal | -3.36 |
|  |  |  | ctx-rh-entorhinal | -3.07 |
|  |  |  | ctx-rh-parahippocampal | -3.07 |
|  |  |  | ctx-rh-fusiform | -2.92 |
|  |  |  | ctx-rh-isthmuscingulate | -2.9 |
|  |  |  | ctx-rh-inferiortemporal | -2.89 |
|  |  |  | ctx-rh-rostralmiddlefrontal | -2.23 |
| Imagination | recall | 1 | Right-Thalamus-Proper | 4.85 |
|  |  |  | ctx-rh-posteriorcingulate | 4.79 |
|  |  |  | Right-Caudate | 4.23 |
|  |  |  | ctx-rh-caudalanteriorcingulate | 4.22 |
|  |  |  | ctx-rh-postcentral | 4.05 |
|  |  |  | ctx-rh-supramarginal | 3.87 |
|  |  |  | ctx-rh-precentral | 3.79 |
|  |  |  | Right-Pallidum | 3.66 |
|  |  |  | ctx-rh-middletemporal | 3.52 |
|  |  |  | ctx-rh-superiorfrontal | 3.47 |
|  |  |  | ctx-rh-superiortemporal | 3.34 |
|  |  |  | Right-Putamen | 3.26 |
|  |  |  | Right-Accumbens-area | 3.24 |
|  |  |  | ctx-rh-transversetemporal | 3.24 |
|  |  |  | ctx-rh-insula | 3.23 |
|  |  |  | ctx-rh-paracentral | 3.06 |
|  |  |  | ctx-lh-entorhinal | 2.58 |
|  |  |  | ctx-rh-caudalmiddlefrontal | 2.58 |
|  |  |  | Right-Amygdala | 2.49 |
|  |  |  | Right-Hippocampus | 2.25 |
|  |  | 2 | ctx-rh-cuneus | -4.62 |
|  |  |  | ctx-rh-pericalcarine | -4.62 |
|  |  |  | ctx-rh-precuneus | -4.6 |
|  |  |  | ctx-rh-isthmuscingulate | -4.41 |
|  |  |  | ctx-rh-lingual | -4.34 |
|  |  |  | ctx-rh-fusiform | -4.02 |
|  |  |  | ctx-rh-lateraloccipital | -3.53 |
|  |  |  | ctx-rh-inferiorparietal | -2.75 |
|  |  |  | ctx-rh-inferiortemporal | -2.66 |
|  |  |  | Right-Hippocampus | -2.29 |
|  |  |  | ctx-rh-middletemporal | -2.23 |
|  | manipulation | 1 | Right-Thalamus-Proper | 4.64 |
|  |  |  | Right-Caudate | 4.21 |
|  |  |  | Right-Pallidum | 3.99 |
|  |  |  | ctx-rh-parsopercularis | 3.39 |
|  |  |  | ctx-rh-precentral | 3.36 |
|  |  |  | Right-Putamen | 3.36 |
|  |  |  | ctx-rh-caudalmiddlefrontal | 3.36 |
|  |  |  | Right-Accumbens-area | 3.34 |
|  |  |  | Right-Amygdala | 3.24 |
|  |  |  | Right-Hippocampus | 3.16 |
|  |  |  | ctx-rh-postcentral | 3.09 |
|  |  |  | ctx-rh-insula | 3.01 |
|  |  |  | ctx-rh-posteriorcingulate | 2.81 |
|  |  |  | ctx-rh-rostralmiddlefrontal | 2.71 |
|  |  |  | ctx-rh-entorhinal | 2.57 |
|  |  |  | ctx-rh-superiorfrontal | 2.55 |
|  |  |  | ctx-rh-supramarginal | 2.46 |
|  |  |  | ctx-rh-caudalanteriorcingulate | 2.11 |
| Third minus first sound | recall | 1 | ctx-rh-medialorbitofrontal | 5.53 |
|  |  |  | ctx-rh-lateralorbitofrontal | 4.8 |
|  |  |  | ctx-rh-parsorbitalis | 4.75 |
|  |  |  | ctx-rh-rostralanteriorcingulate | 4.75 |
|  |  |  | ctx-rh-parstriangularis | 4.36 |
|  |  |  | ctx-rh-rostralmiddlefrontal | 4.34 |
|  |  |  | Right-Accumbens-area | 4.31 |
|  |  |  | ctx-rh-parsopercularis | 4.22 |
|  |  |  | Right-Caudate | 3.96 |
|  |  |  | ctx-rh-insula | 3.58 |
|  |  |  | ctx-rh-superiortemporal | 3.2 |
|  |  |  | Right-Putamen | 3.14 |
|  |  |  | ctx-rh-superiorfrontal | 2.86 |
|  |  |  | ctx-rh-caudalmiddlefrontal | 2.56 |
|  |  |  | ctx-rh-temporalpole | 2.09 |
|  |  | 2 | ctx-rh-inferiortemporal | -6.15 |
|  |  |  | ctx-rh-middletemporal | -5.86 |
|  |  |  | ctx-rh-fusiform | -5.48 |
|  |  |  | ctx-rh-superiortemporal | -5.24 |
|  |  |  | ctx-rh-bankssts | -5.22 |
|  |  |  | ctx-rh-postcentral | -5.01 |
|  |  |  | ctx-rh-supramarginal | -4.34 |
|  |  |  | ctx-rh-parahippocampal | -4.17 |
|  |  |  | Right-Hippocampus | -4.14 |
|  |  |  | ctx-rh-transversetemporal | -3.89 |
|  |  |  | ctx-rh-insula | -3.64 |
|  |  |  | ctx-rh-precentral | -3.51 |
|  |  |  | Right-Putamen | -3.37 |
|  |  |  | Right-Thalamus-Proper | -3.29 |
|  |  |  | Right-Pallidum | -3.25 |
|  |  |  | ctx-rh-entorhinal | -3.1 |
|  |  |  | Right-Amygdala | -2.56 |
|  |  |  | ctx-rh-isthmuscingulate | -2.14 |
|  | manipulation | 1 | ctx-rh-parsorbitalis | 4 |
|  |  |  | ctx-rh-lateralorbitofrontal | 3.89 |
|  |  |  | ctx-rh-parstriangularis | 3.78 |
|  |  |  | ctx-rh-rostralmiddlefrontal | 3.74 |
|  |  |  | Right-Accumbens-area | 3.26 |
|  |  |  | ctx-rh-parsopercularis | 3.18 |
|  |  |  | ctx-rh-rostralanteriorcingulate | 3.06 |
|  |  |  | Right-Caudate | 3.02 |
|  |  |  | ctx-rh-medialorbitofrontal | 3.02 |
|  |  |  | ctx-rh-insula | 2.77 |
|  |  |  | ctx-rh-superiortemporal | 2.51 |
|  |  | 2 | ctx-rh-middletemporal | -5.96 |
|  |  |  | ctx-rh-bankssts | -5.77 |
|  |  |  | ctx-rh-superiortemporal | -5.73 |
|  |  |  | ctx-rh-supramarginal | -5.59 |
|  |  |  | ctx-rh-inferiortemporal | -5.5 |
|  |  |  | Right-Thalamus-Proper | -5.26 |
|  |  |  | Right-Hippocampus | -5.22 |
|  |  |  | Right-Putamen | -4.91 |
|  |  |  | Right-Pallidum | -4.86 |
|  |  |  | ctx-rh-postcentral | -4.72 |
|  |  |  | ctx-rh-transversetemporal | -4.61 |
|  |  |  | ctx-rh-parahippocampal | -4.34 |
|  |  |  | ctx-rh-fusiform | -4.31 |
|  |  |  | Right-Amygdala | -4.3 |
|  |  |  | ctx-rh-insula | -4.06 |
|  |  |  | ctx-rh-precentral | -3.96 |
|  |  |  | ctx-rh-entorhinal | -3.76 |
|  |  |  | ctx-rh-inferiorparietal | -3.49 |
|  |  |  | ctx-rh-posteriorcingulate | -2.8 |
|  |  |  | ctx-rh-paracentral | -2.79 |
|  |  |  | Right-Caudate | -2.74 |
|  |  |  | ctx-rh-isthmuscingulate | -2.26 |
| imagination minus listening | recall | 1 | Right-Thalamus-Proper | 7.56 |
|  |  |  | Right-Caudate | 6.42 |
|  |  |  | ctx-rh-posteriorcingulate | 6.2 |
|  |  |  | Right-Accumbens-area | 5.74 |
|  |  |  | Right-Pallidum | 5.6 |
|  |  |  | ctx-rh-caudalanteriorcingulate | 5.47 |
|  |  |  | ctx-rh-superiorfrontal | 5.42 |
|  |  |  | Right-Putamen | 5.18 |
|  |  |  | ctx-rh-paracentral | 4.91 |
|  |  |  | Right-Amygdala | 4.66 |
|  |  |  | ctx-rh-postcentral | 4.53 |
|  |  |  | ctx-rh-precentral | 4.3 |
|  |  |  | Right-Hippocampus | 4.23 |
|  |  |  | ctx-rh-middletemporal | 4.04 |
|  |  |  | ctx-rh-caudalmiddlefrontal | 4 |
|  |  |  | ctx-rh-superiortemporal | 3.98 |
|  |  |  | ctx-rh-transversetemporal | 3.93 |
|  |  |  | ctx-rh-insula | 3.92 |
|  |  |  | ctx-rh-supramarginal | 3.85 |
|  |  |  | ctx-rh-medialorbitofrontal | 3.74 |
|  |  |  | ctx-rh-parsopercularis | 3.34 |
|  |  |  | ctx-rh-entorhinal | 3.29 |
|  |  |  | ctx-rh-lateralorbitofrontal | 3.1 |
|  |  |  | ctx-rh-rostralmiddlefrontal | 2.57 |
|  |  |  | ctx-rh-inferiortemporal | 2.47 |
|  |  |  | ctx-rh-parahippocampal | 2.46 |
|  |  |  | ctx-rh-isthmuscingulate | 2.45 |
|  |  | 2 | ctx-rh-lingual | -6.44 |
|  |  |  | ctx-rh-isthmuscingulate | -6.13 |
|  |  |  | ctx-rh-precuneus | -5.75 |
|  |  |  | ctx-rh-pericalcarine | -5.25 |
|  |  |  | ctx-rh-cuneus | -5.2 |
|  |  |  | ctx-rh-lateraloccipital | -4.24 |
|  |  |  | ctx-rh-fusiform | -4.14 |
|  |  |  | ctx-rh-inferiortemporal | -3.88 |
|  |  |  | ctx-rh-middletemporal | -3.61 |
|  |  |  | ctx-rh-inferiorparietal | -3.61 |
|  |  |  | ctx-rh-parahippocampal | -3.44 |
|  |  |  | Right-Hippocampus | -3.24 |
|  |  |  | ctx-rh-bankssts | -2.7 |
|  |  |  | ctx-rh-superiorparietal | -2.46 |
|  | manipulation | 1 | Right-Thalamus-Proper | 6.25 |
|  |  |  | Right-Caudate | 4.95 |
|  |  |  | ctx-rh-posteriorcingulate | 4.66 |
|  |  |  | ctx-rh-caudalmiddlefrontal | 4.58 |
|  |  |  | Right-Pallidum | 4.54 |
|  |  |  | ctx-rh-parsopercularis | 4.53 |
|  |  |  | Right-Putamen | 4.44 |
|  |  |  | ctx-rh-superiorfrontal | 4.38 |
|  |  |  | ctx-rh-caudalanteriorcingulate | 4.35 |
|  |  |  | Right-Accumbens-area | 4.3 |
|  |  |  | ctx-rh-precentral | 4.11 |
|  |  |  | ctx-rh-insula | 3.71 |
|  |  |  | Right-Amygdala | 3.29 |
|  |  |  | ctx-rh-paracentral | 3.27 |
|  |  |  | ctx-rh-rostralmiddlefrontal | 3.21 |
|  |  |  | ctx-rh-postcentral | 3.07 |
|  |  |  | Right-Hippocampus | 3.02 |
|  |  |  | ctx-rh-medialorbitofrontal | 2.79 |
|  |  |  | ctx-rh-lateralorbitofrontal | 2.52 |
|  |  |  | ctx-rh-parstriangularis | 2.43 |

**Table S5.** Peak t-value for each region in the **left** hemisphere that overlapped with significant clusters in each of the tested periods and conditions. The Desikan-Killiany parcellation was used to obtain anatomic labels.

| **Period / comparison** | **Condition** | **Cluster No** | **label** | **t value** |
| --- | --- | --- | --- | --- |
| First sound | recall | 1 | ctx-lh-precuneus | 4.46 |
|  |  |  | ctx-lh-bankssts | 4.34 |
|  |  |  | ctx-lh-middletemporal | 4.28 |
|  |  |  | ctx-lh-inferiortemporal | 4.17 |
|  |  |  | ctx-lh-fusiform | 4.15 |
|  |  |  | ctx-lh-cuneus | 4 |
|  |  |  | ctx-lh-inferiorparietal | 3.88 |
|  |  |  | ctx-lh-superiorparietal | 3.87 |
|  |  |  | ctx-lh-supramarginal | 3.57 |
|  |  |  | ctx-lh-superiortemporal | 3.38 |
|  |  |  | ctx-lh-parahippocampal | 3.31 |
|  |  |  | ctx-lh-lingual | 3 |
|  |  |  | ctx-lh-isthmuscingulate | 2.92 |
|  |  |  | Left-Hippocampus | 2.87 |
|  |  |  | ctx-lh-pericalcarine | 2.75 |
|  |  |  | ctx-lh-lateraloccipital | 2.73 |
|  |  | 2 | Left-Accumbens-area | -7.4 |
|  |  |  | Left-Caudate | -7.1 |
|  |  |  | Left-Thalamus-Proper | -6.72 |
|  |  |  | ctx-lh-medialorbitofrontal | -6.36 |
|  |  |  | Left-Pallidum | -6.23 |
|  |  |  | Left-Putamen | -6.15 |
|  |  |  | ctx-lh-rostralanteriorcingulate | -5.97 |
|  |  |  | ctx-lh-lateralorbitofrontal | -5.45 |
|  |  |  | ctx-lh-insula | -5.18 |
|  |  |  | ctx-lh-parsopercularis | -4.99 |
|  |  |  | ctx-lh-parstriangularis | -4.95 |
|  |  |  | ctx-lh-caudalmiddlefrontal | -4.1 |
|  |  |  | ctx-lh-rostralmiddlefrontal | -3.99 |
|  |  |  | Left-Amygdala | -3.83 |
|  |  |  | ctx-lh-posteriorcingulate | -3.67 |
|  |  |  | ctx-lh-precentral | -3.39 |
|  |  |  | ctx-lh-parsorbitalis | -3 |
|  |  |  | Left-Hippocampus | -3 |
|  |  |  | ctx-lh-superiortemporal | -2.95 |
|  |  |  | ctx-lh-inferiortemporal | -2.91 |
|  |  |  | ctx-lh-entorhinal | -2.89 |
|  |  |  | ctx-lh-temporalpole | -2.85 |
|  |  |  | ctx-lh-caudalanteriorcingulate | -2.4 |
|  |  |  | ctx-lh-fusiform | -2.32 |
|  |  |  | ctx-lh-isthmuscingulate | -2.15 |
|  |  |  | ctx-lh-precuneus | -2.06 |
|  | manipulation | 1 | ctx-lh-middletemporal | 5.14 |
|  |  |  | ctx-lh-bankssts | 5.04 |
|  |  |  | ctx-lh-inferiorparietal | 4.86 |
|  |  |  | ctx-lh-inferiortemporal | 4.71 |
|  |  |  | ctx-lh-lateraloccipital | 4.64 |
|  |  |  | ctx-lh-fusiform | 4.62 |
|  |  |  | ctx-lh-lingual | 3.76 |
|  |  |  | ctx-lh-superiortemporal | 3.68 |
|  |  |  | ctx-lh-supramarginal | 3.62 |
|  |  |  | ctx-lh-parahippocampal | 3.42 |
|  |  |  | Left-Hippocampus | 2.93 |
|  |  |  | ctx-lh-superiorparietal | 2.8 |
|  |  |  | ctx-lh-isthmuscingulate | 2.54 |
|  |  |  | ctx-lh-precuneus | 2.49 |
|  |  |  | ctx-lh-cuneus | 2.29 |
|  |  | 3 | Left-Caudate | -5.72 |
|  |  |  | Left-Thalamus-Proper | -5.56 |
|  |  |  | Left-Putamen | -5.45 |
|  |  |  | Left-Accumbens-area | -5.34 |
|  |  |  | Left-Pallidum | -5.17 |
|  |  |  | ctx-lh-insula | -4.95 |
|  |  |  | ctx-lh-parsopercularis | -4.76 |
|  |  |  | ctx-lh-medialorbitofrontal | -4.63 |
|  |  |  | ctx-lh-parstriangularis | -4.49 |
|  |  |  | ctx-lh-rostralanteriorcingulate | -4.36 |
|  |  |  | ctx-lh-caudalmiddlefrontal | -3.84 |
|  |  |  | ctx-lh-lateralorbitofrontal | -3.82 |
|  |  |  | ctx-lh-precentral | -3.78 |
|  |  |  | ctx-lh-rostralmiddlefrontal | -3.38 |
|  |  |  | Left-Amygdala | -3.35 |
|  |  |  | ctx-lh-posteriorcingulate | -3.07 |
|  |  |  | ctx-lh-postcentral | -3.06 |
|  |  |  | ctx-lh-superiortemporal | -2.95 |
|  |  |  | Left-Hippocampus | -2.77 |
|  |  |  | ctx-lh-caudalanteriorcingulate | -2.35 |
|  |  |  | ctx-lh-entorhinal | -2.3 |
|  |  |  | ctx-lh-temporalpole | -2.28 |
|  |  |  | ctx-lh-supramarginal | -2.11 |
| Second sound | recall | 1 | ctx-lh-fusiform | 4.73 |
|  |  |  | ctx-lh-lingual | 4.26 |
|  |  |  | ctx-lh-parahippocampal | 4.21 |
|  |  |  | ctx-lh-isthmuscingulate | 4.06 |
|  |  |  | ctx-lh-precuneus | 3.76 |
|  |  |  | Left-Hippocampus | 3.75 |
|  |  |  | ctx-lh-inferiortemporal | 3.31 |
|  |  |  | ctx-lh-pericalcarine | 3.21 |
|  |  |  | ctx-lh-cuneus | 3.15 |
|  |  |  | ctx-lh-superiorparietal | 3.15 |
|  |  |  | ctx-lh-middletemporal | 2.93 |
|  |  |  | ctx-lh-bankssts | 2.69 |
|  |  |  | ctx-lh-superiortemporal | 2.65 |
|  |  |  | ctx-lh-supramarginal | 2.55 |
|  |  | 2 | Left-Thalamus-Proper | -4.58 |
|  |  |  | ctx-lh-posteriorcingulate | -4.03 |
|  |  |  | Left-Caudate | -3.54 |
|  |  |  | Left-Accumbens-area | -3.28 |
|  |  |  | ctx-lh-caudalanteriorcingulate | -2.81 |
|  |  |  | Left-Pallidum | -2.68 |
|  |  |  | ctx-lh-isthmuscingulate | -2.54 |
|  |  |  | ctx-lh-medialorbitofrontal | -2.33 |
|  |  |  | Left-Putamen | -2.21 |
|  |  |  | ctx-lh-rostralanteriorcingulate | -2.18 |
|  | manipulation | 1 | ctx-lh-precentral | -4.56 |
|  |  |  | ctx-lh-caudalmiddlefrontal | -3.77 |
|  |  |  | ctx-lh-postcentral | -3.58 |
|  |  |  | ctx-lh-caudalanteriorcingulate | -3.51 |
|  |  |  | Left-Thalamus-Proper | -3.43 |
|  |  |  | ctx-lh-posteriorcingulate | -3.36 |
|  |  |  | ctx-lh-superiorfrontal | -3.34 |
|  |  |  | Left-Pallidum | -2.94 |
|  |  |  | ctx-lh-inferiortemporal | -2.82 |
|  |  |  | ctx-lh-paracentral | -2.71 |
|  |  |  | Left-Caudate | -2.63 |
|  |  |  | Left-Amygdala | -2.61 |
|  |  |  | Left-Putamen | -2.59 |
|  |  |  | Left-Hippocampus | -2.52 |
|  |  |  | ctx-lh-superiortemporal | -2.37 |
|  |  |  | ctx-lh-middletemporal | -2.1 |
|  |  |  | ctx-lh-fusiform | -2.09 |
| Third sound | recall | 1 | ctx-lh-lingual | 3.08 |
|  |  |  | ctx-lh-precuneus | 2.76 |
|  |  |  | ctx-lh-cuneus | 2.71 |
|  |  |  | ctx-lh-lateraloccipital | 2.69 |
|  |  |  | ctx-lh-pericalcarine | 2.64 |
|  |  |  | ctx-lh-isthmuscingulate | 2.5 |
|  |  | 2 | Left-Thalamus-Proper | -5 |
|  |  |  | ctx-lh-superiortemporal | -4.83 |
|  |  |  | ctx-lh-transversetemporal | -4.7 |
|  |  |  | ctx-lh-postcentral | -4.6 |
|  |  |  | ctx-lh-supramarginal | -4.4 |
|  |  |  | ctx-lh-bankssts | -4.35 |
|  |  |  | ctx-lh-middletemporal | -4.21 |
|  |  |  | ctx-lh-insula | -3.54 |
|  |  |  | ctx-lh-inferiortemporal | -3.51 |
|  |  |  | ctx-lh-precentral | -3.35 |
|  |  |  | Left-Caudate | -3.12 |
|  |  |  | Left-Putamen | -2.71 |
|  |  |  | ctx-lh-inferiorparietal | -2.7 |
|  |  |  | ctx-lh-fusiform | -2.59 |
|  |  |  | ctx-lh-posteriorcingulate | -2.58 |
|  |  |  | Left-Hippocampus | -2.41 |
|  |  |  | Left-Pallidum | -2.21 |
|  |  |  | ctx-lh-parahippocampal | -2.15 |
|  | manipulation | 1 | ctx-lh-superiortemporal | -4.66 |
|  |  |  | ctx-lh-supramarginal | -4.54 |
|  |  |  | ctx-lh-bankssts | -4.23 |
|  |  |  | Left-Thalamus-Proper | -4.02 |
|  |  |  | ctx-lh-middletemporal | -3.98 |
|  |  |  | ctx-lh-postcentral | -3.96 |
|  |  |  | ctx-lh-precentral | -3.83 |
|  |  |  | ctx-lh-transversetemporal | -3.33 |
|  |  |  | Left-Hippocampus | -2.89 |
|  |  |  | Left-Amygdala | -2.85 |
|  |  |  | ctx-lh-inferiortemporal | -2.79 |
|  |  |  | ctx-lh-parahippocampal | -2.66 |
|  |  |  | ctx-lh-insula | -2.61 |
|  |  |  | ctx-lh-entorhinal | -2.58 |
|  |  |  | Left-Caudate | -2.51 |
|  |  |  | ctx-lh-fusiform | -2.43 |
|  |  |  | Left-Pallidum | -2.42 |
|  |  |  | Left-Putamen | -2.27 |
| Imagination | recall | 1 | Left-Thalamus-Proper | 4.95 |
|  |  |  | ctx-lh-posteriorcingulate | 4.93 |
|  |  |  | Left-Pallidum | 4.14 |
|  |  |  | ctx-lh-caudalanteriorcingulate | 4.05 |
|  |  |  | Left-Caudate | 4 |
|  |  |  | Left-Putamen | 3.51 |
|  |  |  | Left-Accumbens-area | 3.41 |
|  |  |  | ctx-lh-precentral | 3.15 |
|  |  |  | Left-Amygdala | 3.07 |
|  |  |  | Left-Hippocampus | 3.04 |
|  |  |  | ctx-lh-superiorfrontal | 2.91 |
|  |  |  | ctx-lh-insula | 2.68 |
|  |  |  | ctx-lh-caudalmiddlefrontal | 2.63 |
|  |  |  | ctx-lh-entorhinal | 2.58 |
|  |  |  | ctx-lh-rostralmiddlefrontal | 2.51 |
|  |  |  | ctx-lh-paracentral | 2.44 |
|  |  |  | ctx-lh-superiortemporal | 2.4 |
|  |  |  | ctx-lh-postcentral | 2.14 |
|  |  | 2 | ctx-lh-precuneus | -4.37 |
|  |  |  | ctx-lh-cuneus | -4.34 |
|  |  |  | ctx-lh-isthmuscingulate | -4.32 |
|  |  |  | ctx-lh-pericalcarine | -3.95 |
|  |  |  | ctx-lh-lingual | -3.49 |
|  |  |  | ctx-lh-middletemporal | -3.37 |
|  |  |  | ctx-lh-lateraloccipital | -3.37 |
|  |  |  | ctx-lh-inferiorparietal | -3.37 |
|  |  |  | ctx-lh-bankssts | -3.3 |
|  |  |  | ctx-lh-superiorparietal | -3.09 |
|  |  |  | ctx-lh-superiortemporal | -2.95 |
|  |  |  | ctx-lh-supramarginal | -2.93 |
|  |  |  | Left-Hippocampus | -2.78 |
|  |  |  | ctx-lh-inferiortemporal | -2.74 |
|  |  |  | ctx-lh-parahippocampal | -2.32 |
|  |  |  | ctx-lh-fusiform | -2.22 |
|  | manipulation | 1 | Left-Thalamus-Proper | 3.91 |
|  |  |  | ctx-lh-entorhinal | 2.37 |
|  |  |  | ctx-lh-parahippocampal | 2.37 |
|  |  |  | Left-Hippocampus | 2.34 |
|  |  |  | Left-Amygdala | 2.24 |
|  | difference | 1 | ctx-lh-parsopercularis | -3.34 |
|  |  |  | ctx-lh-caudalmiddlefrontal | -3.05 |
|  |  |  | ctx-lh-precentral | -2.88 |
|  |  |  | ctx-lh-insula | -2.8 |
|  |  |  | ctx-lh-rostralmiddlefrontal | -2.78 |
|  |  |  | ctx-lh-parstriangularis | -2.76 |
|  |  |  | ctx-lh-lateralorbitofrontal | -2.76 |
|  |  |  | Left-Thalamus-Proper | -2.71 |
|  |  |  | Left-Putamen | -2.66 |
|  |  |  | ctx-lh-superiorfrontal | -2.65 |
|  |  |  | Left-Caudate | -2.58 |
|  |  |  | ctx-lh-parsorbitalis | -2.49 |
|  |  |  | Left-Pallidum | -2.48 |
|  |  |  | ctx-lh-posteriorcingulate | -2.1 |
|  |  |  | ctx-lh-superiortemporal | -2.05 |
| Third minus first sound | recall | 1 | ctx-lh-medialorbitofrontal | 5.11 |
|  |  |  | ctx-lh-rostralanteriorcingulate | 4.82 |
|  |  |  | Left-Accumbens-area | 4.6 |
|  |  |  | ctx-lh-lateralorbitofrontal | 4.52 |
|  |  |  | Left-Putamen | 4.12 |
|  |  |  | Left-Caudate | 4.09 |
|  |  |  | ctx-lh-insula | 3.69 |
|  |  |  | Left-Pallidum | 3.44 |
|  |  |  | ctx-lh-parstriangularis | 3.23 |
|  |  |  | ctx-lh-parsorbitalis | 3.16 |
|  |  |  | ctx-lh-parsopercularis | 3.08 |
|  |  |  | ctx-lh-superiortemporal | 3.07 |
|  |  |  | ctx-lh-rostralmiddlefrontal | 2.96 |
|  |  |  | ctx-lh-superiorfrontal | 2.89 |
|  |  |  | ctx-lh-temporalpole | 2.72 |
|  |  |  | ctx-lh-caudalmiddlefrontal | 2.4 |
|  |  |  | Left-Thalamus-Proper | 2.2 |
|  |  | 2 | Left-Thalamus-Proper | -2.31 |
|  |  | 3 | ctx-lh-middletemporal | -4.78 |
|  |  |  | ctx-lh-bankssts | -4.65 |
|  |  |  | ctx-lh-superiortemporal | -4.53 |
|  |  |  | ctx-lh-inferiorparietal | -4.41 |
|  |  |  | ctx-lh-inferiortemporal | -4.28 |
|  |  |  | ctx-lh-supramarginal | -3.99 |
|  |  |  | ctx-lh-fusiform | -3.95 |
|  |  |  | ctx-lh-postcentral | -3.44 |
|  |  |  | ctx-lh-transversetemporal | -3.32 |
|  |  |  | Left-Hippocampus | -3.01 |
|  |  |  | ctx-lh-parahippocampal | -2.97 |
|  |  |  | ctx-lh-insula | -2.55 |
|  | manipulation | 1 | Left-Caudate | 3.92 |
|  |  |  | ctx-lh-insula | 3.88 |
|  |  |  | ctx-lh-parstriangularis | 3.78 |
|  |  |  | Left-Putamen | 3.57 |
|  |  |  | ctx-lh-parsopercularis | 3.53 |
|  |  |  | ctx-lh-medialorbitofrontal | 3.42 |
|  |  |  | ctx-lh-lateralorbitofrontal | 3.31 |
|  |  |  | ctx-lh-rostralanteriorcingulate | 3.3 |
|  |  |  | Left-Accumbens-area | 3.24 |
|  |  |  | ctx-lh-superiorfrontal | 2.92 |
|  |  |  | ctx-lh-precentral | 2.88 |
|  |  |  | Left-Pallidum | 2.82 |
|  |  |  | ctx-lh-parsorbitalis | 2.72 |
|  |  |  | ctx-lh-rostralmiddlefrontal | 2.62 |
|  |  |  | ctx-lh-postcentral | 2.54 |
|  |  |  | ctx-lh-superiortemporal | 2.47 |
|  |  |  | ctx-lh-caudalmiddlefrontal | 2.37 |
|  |  | 2 | ctx-lh-middletemporal | -5.19 |
|  |  |  | ctx-lh-bankssts | -5.11 |
|  |  |  | ctx-lh-inferiortemporal | -4.65 |
|  |  |  | ctx-lh-superiortemporal | -4.45 |
|  |  |  | ctx-lh-inferiorparietal | -3.99 |
|  |  |  | ctx-lh-supramarginal | -3.77 |
|  |  |  | ctx-lh-fusiform | -3.5 |
|  |  |  | ctx-lh-parahippocampal | -3.24 |
|  |  |  | Left-Hippocampus | -3.19 |
|  |  |  | ctx-lh-transversetemporal | -2.87 |
|  |  |  | ctx-lh-lingual | -2.82 |
|  |  |  | ctx-lh-isthmuscingulate | -2.23 |
| imagination minus listening | recall | 1 | Left-Thalamus-Proper | 8.23 |
|  |  |  | Left-Caudate | 6.94 |
|  |  |  | Left-Pallidum | 6.24 |
|  |  |  | Left-Accumbens-area | 6.22 |
|  |  |  | ctx-lh-posteriorcingulate | 6.18 |
|  |  |  | Left-Putamen | 5.14 |
|  |  |  | ctx-lh-caudalanteriorcingulate | 4.64 |
|  |  |  | Left-Amygdala | 4.44 |
|  |  |  | Left-Hippocampus | 4.17 |
|  |  |  | ctx-lh-precentral | 3.93 |
|  |  |  | ctx-lh-medialorbitofrontal | 3.52 |
|  |  |  | ctx-lh-isthmuscingulate | 3.38 |
|  |  |  | ctx-lh-inferiortemporal | 3.36 |
|  |  |  | ctx-lh-postcentral | 3.31 |
|  |  |  | ctx-lh-entorhinal | 3.12 |
|  |  |  | ctx-lh-caudalmiddlefrontal | 2.91 |
|  |  |  | ctx-lh-insula | 2.89 |
|  |  |  | ctx-lh-superiortemporal | 2.86 |
|  |  |  | ctx-lh-rostralanteriorcingulate | 2.81 |
|  |  |  | ctx-lh-transversetemporal | 2.79 |
|  |  |  | ctx-lh-fusiform | 2.72 |
|  |  |  | ctx-lh-lateralorbitofrontal | 2.65 |
|  |  |  | ctx-lh-rostralmiddlefrontal | 2.65 |
|  |  |  | ctx-lh-superiorfrontal | 2.6 |
|  |  |  | ctx-lh-parsopercularis | 2.22 |
|  |  |  | ctx-lh-parstriangularis | 2.2 |
|  |  |  | ctx-lh-supramarginal | 2.09 |
|  |  | 2 | ctx-lh-precuneus | -6.5 |
|  |  |  | ctx-lh-cuneus | -5.67 |
|  |  |  | ctx-lh-superiorparietal | -5.3 |
|  |  |  | ctx-lh-isthmuscingulate | -5.22 |
|  |  |  | ctx-lh-fusiform | -5.01 |
|  |  |  | ctx-lh-pericalcarine | -4.99 |
|  |  |  | ctx-lh-middletemporal | -4.37 |
|  |  |  | ctx-lh-bankssts | -4.32 |
|  |  |  | ctx-lh-inferiorparietal | -4.29 |
|  |  |  | ctx-lh-lingual | -4.19 |
|  |  |  | ctx-lh-inferiortemporal | -4.17 |
|  |  |  | ctx-lh-supramarginal | -3.8 |
|  |  |  | ctx-lh-parahippocampal | -3.7 |
|  |  |  | ctx-lh-lateraloccipital | -3.6 |
|  |  |  | ctx-lh-superiortemporal | -3.09 |
|  |  |  | Left-Hippocampus | -2.6 |
|  | manipulation | 1 | Left-Thalamus-Proper | 6.52 |
|  |  |  | Left-Pallidum | 4.53 |
|  |  |  | Left-Accumbens-area | 4.5 |
|  |  |  | Left-Caudate | 4.14 |
|  |  |  | Left-Amygdala | 3.71 |
|  |  |  | Left-Putamen | 3.68 |
|  |  |  | Left-Hippocampus | 3.41 |
|  |  |  | ctx-lh-posteriorcingulate | 3.07 |
|  |  |  | ctx-lh-superiorfrontal | 2.93 |
|  |  |  | ctx-lh-entorhinal | 2.89 |
|  |  |  | ctx-lh-paracentral | 2.82 |
|  |  |  | ctx-lh-middletemporal | 2.74 |
|  |  |  | ctx-lh-parahippocampal | 2.67 |
|  |  |  | ctx-lh-fusiform | 2.66 |
|  |  |  | ctx-lh-insula | 2.63 |
|  |  |  | ctx-lh-superiortemporal | 2.61 |
|  |  |  | ctx-lh-transversetemporal | 2.58 |
|  |  |  | ctx-lh-medialorbitofrontal | 2.57 |
|  |  |  | ctx-lh-caudalanteriorcingulate | 2.39 |
